# A post-processing algorithm for building longitudinal medication dose data from extracted medication information using natural language processing from electronic health records

**DOI:** 10.1101/775015

**Authors:** Elizabeth McNeer, Cole Beck, Hannah L. Weeks, Michael L. Williams, Nathan T. James, Leena Choi

## Abstract

**Objective:** We developed a post-processing algorithm to convert raw natural language processing (NLP) output from electronic health records (EHRs) into a usable format for analysis. This algorithm was specifically developed for creating datasets for use in medication-based studies.

**Materials and Methods:** The algorithm was developed using output from two NLP systems, MedXN and medExtractR. We extracted medication information from deidentified clinical notes from Vanderbilt’s EHR system for two medications, tacrolimus and lamotrigine. The algorithm consists of two parts. Part I parses the raw NLP output and connects entities together. Part II removes redundancies and calculates dose intake and daily dose. We evaluated each part by comparing to human-determined gold standards that were generated using approximately 300 records from 10 subjects for each medication and each NLP system.

**Results:** The algorithm performed well. For MedXN, the F-measures were at or above 0.99 for Part I and at or above 0.97 for Part II. For medExtractR, the F-measures for Part I were 1.00 and for Part II they were at or above 0.98.

**Discussion:** Our post-processing algorithm was developed separately from an NLP system, making it easier to modify and generalize to other systems. It performed well to convert NLP output to analyzable data, but it cannot perform well in certain cases, such as when incorrect information is extracted by the NLP system.

**Conclusion:** Our post-processing algorithm provides a way to convert raw NLP output to a form that is useful for medication-based studies, leading to more opportunities to use EHR data for diverse studies.

## INTRODUCTION

Natural language processing (NLP) systems are very useful in a variety of settings. The development of NLP systems for extracting information from electronic health records (EHRs) has led to great opportunities for performing diverse research by providing critical pieces of data. Many NLP systems have been developed specifically for clinical research. For example, cTAKES,[1] MetaMap,[2] and MedLEE[3] are NLP systems for general purpose extraction of clinical information. In early work, Evans et al. used the CLARIT system to extract medication names and dosage information from clinical narrative text.[4] Their system extracted medication dosage information with about 80% accuracy. More recently, several NLP systems such as MedEx[5], CLAMP,[6] MedXN,[7] and medExtractR,[8] have been developed specifically for extracting medication information from EHRs.

Although NLP systems are useful to extract medication information from unstructured text, the raw output from some of these systems is not directly usable and requires a post-processing step to convert the extracted information into an appropriate data form for further analysis depending on the research goal. For example, with medication extraction, the entities (or attributes) of the raw output from NLP systems should be associated with corresponding medication names to make dose data, such as dose given intake or daily dose, and then this data can be used for further analysis in diverse medication-based studies. Some NLP systems have a built-in post-processing step as a part of the system. For example, the final step in the system built by Patrick et al. for the 2009 i2b2 medication extraction challenge was equipped with a medication entry generator for assembling medication events based on the relationships between components established in previous steps.[9] Also, the Lancet system developed by Li *et al.* included a supervised machine learning classifier that attempted to associate a medication name with the correct entities.[10] MedEx[5] and CLAMP[6] also include a post-processing step that connects entities with the associated drug name in a data structure, but they may not provide complete dose data such as dose given intake or daily dose that are often required in many medication-based studies.

In addition, to the best of our knowledge, the validation of post-processing algorithms within these NLP systems has not been reported separately from the overall evaluation of the system in the literature, although they may be unofficially validated during the development phase. After extracting information from unstructured text using an NLP, building useable data can be especially challenging when several extracted entities should be paired to define a phenotype or quantity of interest. This is especially true for making dose data from extracted drug entities. Correctly connecting entities with both drug name and with each other to obtain medication dose (e.g., dose given intake or daily dose) can be challenging and error prone if the prescription pattern is complex. Developing and validating a post-processing algorithm separately from the main NLP extraction system allows for easier identification of error sources and more efficient improvement of relevant parts of the system.

We separately evaluated medication extraction for four NLP systems in our previous work.[8] Out of these four tested NLP systems, we selected the two best-performing systems, MedXN[7] and medExtractR,[8] to develop an algorithm that processes the raw output. The post-processing algorithm we developed is divided into two parts. Part I parses the raw output from the NLP system and connects entities together. Part II removes redundant data entries anchored at either note or date identifiers and calculates dose given intake and daily dose. These dose data are crucial measurements of drug exposure for many medication-based studies including population pharmacokinetic and pharmacodynamic or pharmacogenomic studies, which are often interested in determining drug exposure and response (or risk) relationships. As such, our post-processing algorithm was implemented in a recently developed system for population pharmacokinetic and pharmacodynamic studies using EHRs.[11] This system provides more precise dosing information for diverse medication-based studies, equipped with many modules including data processing modules. Our algorithm is the key component of one of these modules called Pro-Med-NLP, which was only schematically presented with the main functionality. Here we present detailed description of the algorithm along with the algorithm validation using the gold standard.

## METHODS

### Data source

To develop a post-processing algorithm, we used two medications, tacrolimus and lamotrigine, whose prescription patterns vary from simple to complicated. For each medication, we defined a patient cohort separately using patient records in a de-identified database of clinical records derived from Vanderbilt’s EHR system. For tacrolimus data, we used the same cohort (n=466) used in previous studies,[12] who were treated with tacrolimus after renal transplant. For lamotrigine data, we first identified patient records with ‘lamotrigine’ or ‘Lamictal’ (brand name of lamotrigine) and an ICD-9-CM or ICD-10-CM (The International Classification of Diseases, Ninth and Tenth Revision, Clinical Modification) billing code for epilepsy before October 3, 2017. We further refined the cohort by selecting patients who had their first lamotrigine level between 18 and 70 years of age and at least 3 drug levels and 3 doses, which yielded the final cohort of 305 subjects. For each subject of each cohort, we identified all clinical notes generated on the same dates when drug concentration laboratory values were available, from which medication dosing information was extracted using both MedXN and medExtractR.

### Medication entity extraction using existing NLP systems

#### MedXN

The MedXN (Medication Extraction and Normalization) system was designed to extract medication information from clinical notes and convert it into an RxNorm concept unique identifier (RxCUI) for the purpose of medication reconciliation.[7] This system identifies medication names and entities, such as dosage, strength, and frequency, in clinical notes using the RxNorm dictionary and regular expressions. The entities associated with a medication name are formatted in the RxNorm standard order and normalized to a specific RxCUI.

#### medExtractR

MedExtractR is a medication extraction algorithm developed using the R programming language, which uses a more targeted approach for specific drugs of interest.[8] Given a list of drug names to search for, medExtractR creates a search window around each identified drug mention, within which related drug entities are extracted. Some drug entities are identified and extracted based on matching expressions in manually curated entity-specific dictionaries, including frequency, intake time, and dose change (keywords to indicate that a regimen may not be current). For the remaining entities, including strength, dose amount (e.g., number of pills to take), dose (i.e., dose given intake), and time of last dose, regular expressions are used to identify common patterns for how these entities are written. Function arguments can be used to optimize drug entity extraction for a given set of clinical notes and drug of interest. Details of medExtractR can be found in Weeks *et al.*[8]

### Description of post-processing algorithms

Although MedXN provides individual drug entities that are linked to specific drug name mentions, the entities are not paired to provide complete dosing information such as dose given intake or daily dose. MedExtractR also provides individual drug entities, but they are not linked to a drug name mention, so they should be linked to a drug name mention first before being paired. In the context of dose-related studies, the output from these systems can only be analyzed after being combined into complete drug regimens and computing dose quantities. A simple post-processing method may form all possible combinations of extracted entities. Consider a drug mention with two strengths, two dose amounts and two frequencies. Applying this simple method would result in eight combinations of strengths, dose amounts, and frequencies, most of which are incorrect pairings; hence, we developed a more sophisticated post-processing algorithm to identify appropriate pairings of medication entities.

#### Part I

Part I of the algorithm processes the raw NLP output and pairs the parsed entities, as illustrated in **Figure 1**. The algorithm begins with a single file that is the raw output from an NLP system, from which a drug name and its entities are isolated and converted to a standardized form [**Step 1**]. The standardized form, structured similar to MedXN output, includes a row for each drug mention and columns for all entities anchored to that drug mention. The entities include strength, dose amount, route, frequency, duration, dose, and dose change. The dose change and dose entities are only applicable to medExtractR (for details see Weeks *et al.*[8]). Note that examples provided in figures and tables only include some of these entities for simplicity. Next, any records with irrelevant drug names (i.e., names of drugs which are not part of the study) are removed [**Step 2**]. For records with at most one value for each entity of a specific drug mention, the pairing part of the algorithm is not called **[Step 3a]**. For all other records, the pairing algorithm is called **[Step 3b]**, and groups are formed anchored to the closest preceding strength **[Step 4]**. Each group consists of one strength and any subsequent entities that occur before the next strength. Any entities before the first strength in the mention form a group (called group 0) which is assigned a missing value (NA) for strength. If all entities in a group have at most one value, the strength is paired with all entities in that group **[Step 5a]**. If any entity in a group has multiple values, all possible paths are searched to find the path with the minimum cost (for more details of cost calculation, see below), where a “path” represents a complete combination among available entities (e.g., drug name with a strength, dose amount, and frequency) including an assigned value of NA for any entity that is missing **[Step 5b]**. Finally, if there is a unique, standardized value for route (e.g., orally) within a drug mention, that route is added to all pairings for that drug mention **[Step 6]**.

**Figure 1.**
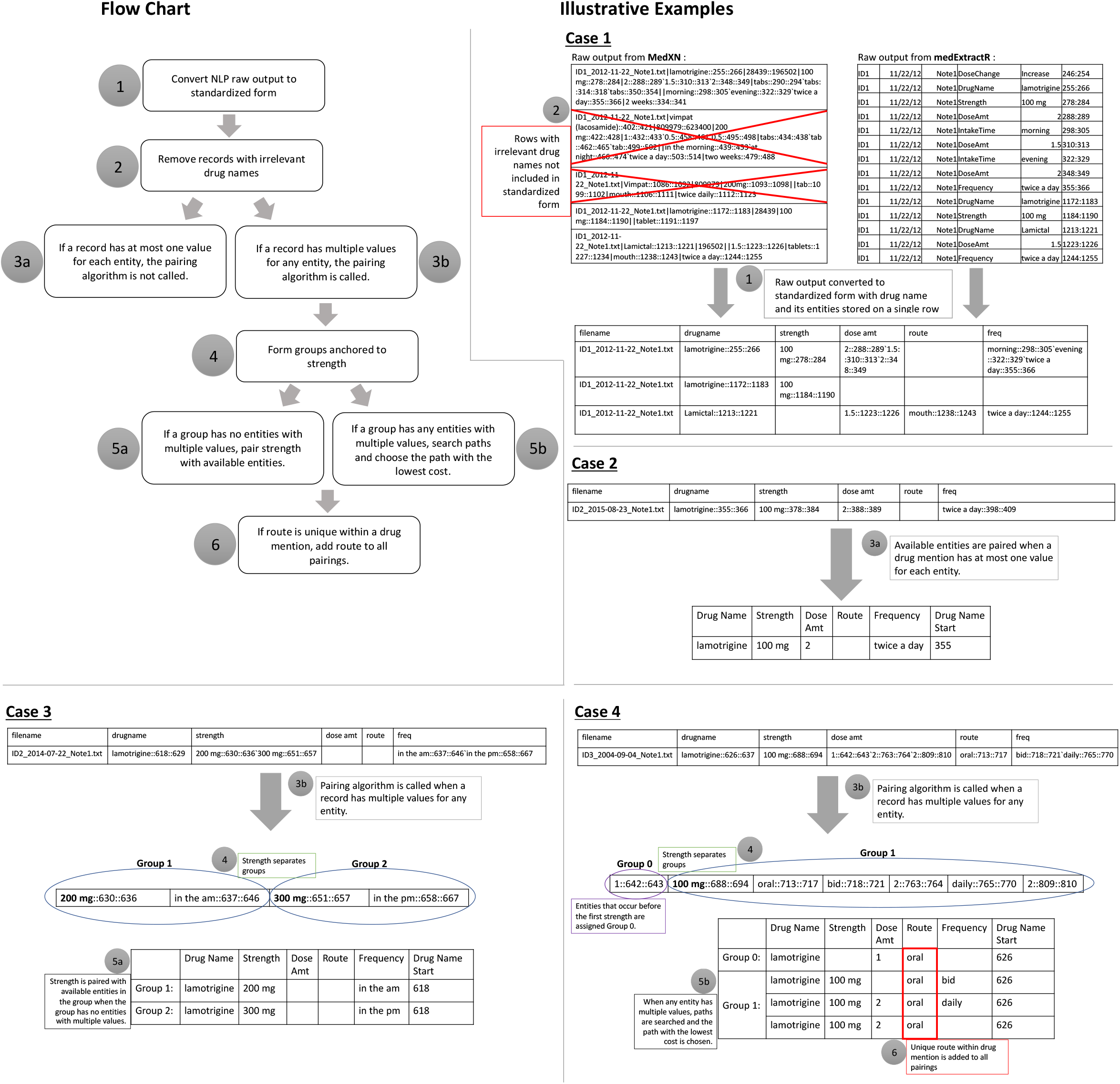
Diagram of Part I of the post-processing algorithm. Part I consists of six main steps. Case 1 illustrates steps 1 and 2, converting the NLP raw output to a standardized form and removing irrelevant drug names. Case 2 illustrates a situation where each entity has at most one value and the pairing algorithm is not called. Case 3 and Case 4 show how groups are formed in step 4. Case 3 illustrates a simple example of forming pairings when there is only one value for each entity. Case 4 shows a more complicated example when at least one entity has multiple values and paths are searched to find the optimal path.

For the cost calculation when searching possible paths in Step 5, we used the distance between entities for each path since we believe the gaps should give insight into which entities are related to each other (i.e., shorter gaps between entities indicates higher likelihood they are related). We considered seven methods for calculating cost, which are described in **Table S1** and illustrated using an example in **Figure S1** of **Supplementary Material**. Among all of the methods, we found that the minimum distance between entity begin positions (minEntBeg) and the minimum distance between entity end position and next entity begin position (minEntEnd) methods performed best. We set the default method to be minEntEnd. The cost calculation for each possible path is illustrated in **Figure S2** of **Supplementary Material** using an example with the default cost method.

#### Part II

Given the output from Part I, Part II of the algorithm standardizes each entity, imputes appropriate missing entities, calculates dose given intake and daily dose, and removes redundant data entries anchored at note or date level for a given patient. An overview of steps along with some examples are presented in **Figure 2**. First, character entities (e.g., strength, dose amount, and frequency) are converted to numeric values where applicable [**Step 1**]. As part of this process, frequency and route values are first standardized as strings. For example, the frequencies “twice a day”, “twice daily”, “bid”, etc. are all standardized to the string “bid”, and routes “oral”, “mouth”, “po”, etc. are all standardized to the string “orally”. Frequencies that give a time of day (e.g., “qam”, “at noon”, “in the evening”) are assigned a numeric value of 1, and a new entity, “intaketime”, is added to record this information using the standardized strings “am”, “noon”, and “pm”. Rows that include only drug name or only drug name and route are removed [**Step 2**]. If there are drug name changes in adjacent rows within the same note (e.g., lamotrigine to Lamictal), these rows are collapsed into one row if there are no conflicts [**Step 3**]. This usually happens when a phrase such as “lamotrigine 100mg (also known as Lamictal) 3 tablets bid” is present in the original note, resulting in two rows of output for the same drug mention. In this example, one row has a drug name of lamotrigine, a strength of 100 and no dose amount or frequency while the next row has a drug name of Lamictal, a dose amount of 3 and a frequency of 2 with no strength. These two rows are combined, yielding a single row with a strength of 100, a dose amount of 3, and a frequency of 2, which provides complete dosing information.

**Figure 2.**
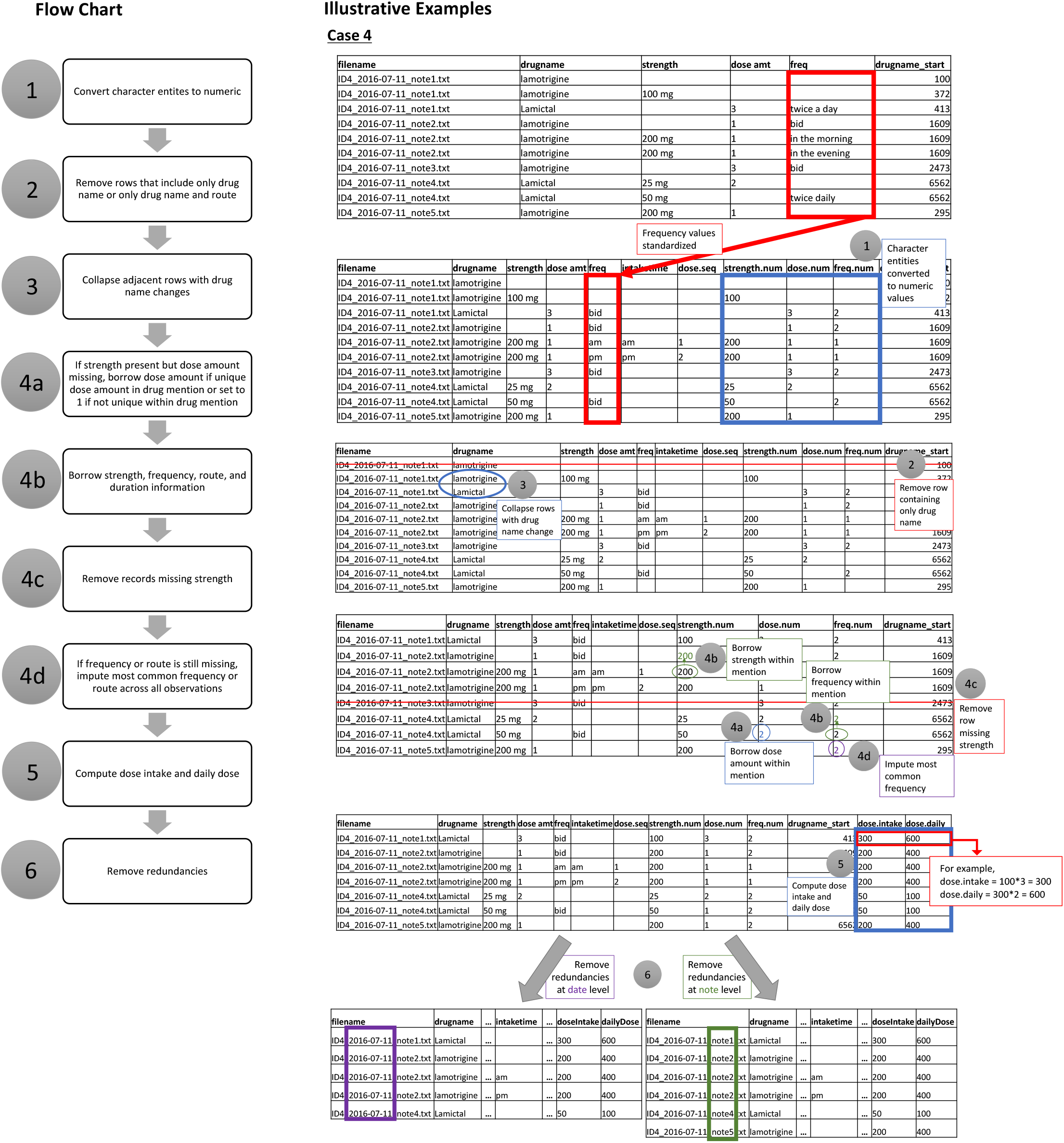
Diagram of Part II of the post-processing algorithm. Part II consists of six main steps. Case 4 illustrates examples of each of these steps. The final output of Part II is two datasets, one with redundancies removed at the date level and one with redundancies removed at the note level.

In the next step, a set of rules are used to impute missing values **[Step 4]**. For example, a missing entity could be imputed by borrowing a unique value within the same drug mention or note or using a global mode. The order in which we apply the rules varies by entity, which is schematically presented in **Figure 3**. Specific examples of precedence rules using the route entity are demonstrated in **Table 1**. In Example 1, this note has a unique route of orally, so this route was imputed in the highlighted and italicized cells where route was originally missing. The note in Example 2 did not have a unique route, so the most common route across all observations for that medication (i.e., orally for tacrolimus) was imputed in the highlighted and italicized cells. When a drug mention is associated with an intake time (e.g., am, noon, pm), it is considered to be part of a dose sequence if at least two of the rows have intake times and the rows are ordered [am, pm] or [am, noon, pm]. Intake time information can be imputed when the observation may be part of a dose sequence. For example, if an observation is missing frequency but the previous observation has an intake time of “am”, then a frequency of “pm” may be used. After applying imputation, any records that are still missing strength are removed. If dose amount is still missing, it is imputed as 1.

**Figure 3.**
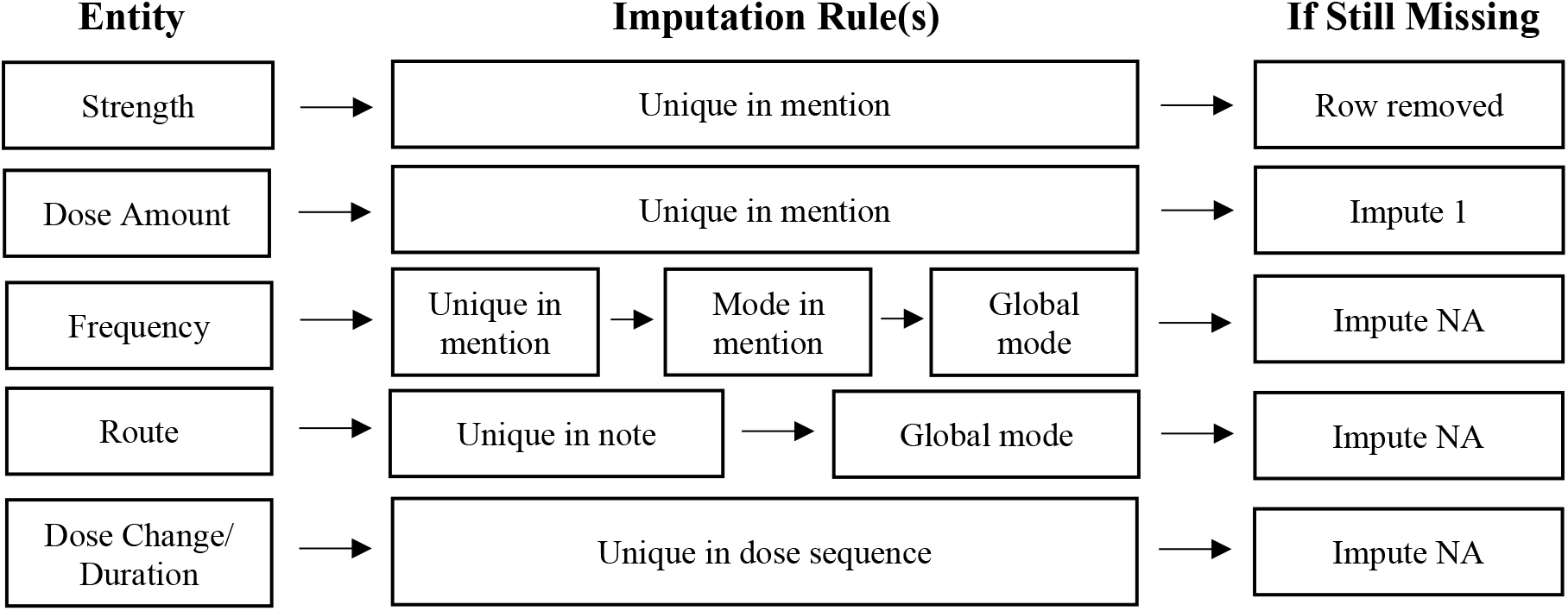
Precedence of rules for imputing missing values in Part I algorithm. The Imputation Rule(s) column shows the rules that are used for imputing missing values for each entity. The rules are ordered from left to right by precedence. For example, if route is missing, the algorithm will first look for a unique route in the note to impute, and then will impute the most common route across all notes if a unique route doesn’t exist in the note. If a value is not imputed, one of three things will happen, as shown in the If Still Missing column.

**Table 1.** Examples of precedence rules for imputing missing values for route.

|  | Filename | Drug Name | Strength | Dose Amount | Route | Frequency | Intake Time | Drug Name Start |
| --- | --- | --- | --- | --- | --- | --- | --- | --- |
| **Example 1** | ID1_2015-04-30_note1.txt | lamotrigine | 100 | 2 | orally | 1 | am | 536 |
|  | ID1_2015-04-30_note1.txt | lamotrigine | 100 | 2 | orally | 1 | pm | 536 |
|  | ID1_2015-04-30_note1.txt | lamotrigine | 200 | 1 | *orally* | 1 | am | 936 |
|  | ID1_2015-04-30_note1.txt | lamotrigine | 300 | 1 | *orally* | 1 | pm | 936 |
| **Example 2** | ID2_2006-05-18_note1.txt | Prograf | 1 | 3 | *orally* | 2 |  | 506 |
|  | ID2_2006-05-18_note1.txt | Prograf | 1 | 3 | *orally* | 2 |  | 506 |

Next, the dose given intake is computed by multiplying strength by dose amount. If there is an intake time, the daily dose is calculated by adding the dose given intakes at each of the intake times. If an intake time is not available, the daily dose is computed by multiplying dose given intake by frequency [**Step 5**]. Finally, any redundancies are removed at date level and note level separately, yielding two datasets [**Step 6**]. Rows were considered redundant if they had the same dose intake, daily dose, dose change, route, and duration, except if one row was part of a dose sequence and the other row was not.

### Generation of training and test sets

We generated training and test sets separately for tacrolimus and lamotrigine from medication entities extracted by each NLP system (MedXN and medExtractR). Each dataset for each drug included approximately 300 clinical notes from 10 patients.

### Making gold standard datasets

We generated three gold standards per dataset to evaluate each part of the post-processing algorithm. Each gold standard dataset was manually generated to make the intended output for each part of the algorithm. That means that we generated “Gold Standard I” as the output of Part I if the six steps of the Part I algorithm were correctly performed. Then, we generated “Gold Standard II–Date” and “Gold Standard II–Note” as the output data of Part II applied to Gold Standard I at the date and note level, respectively, if the six steps of the Part II algorithm were correctly performed. These were generated for each of the training and test sets and each of the medications (i.e., tacrolimus and lamotrigine), yielding a total of 12 gold standard datasets for each NLP system. For each gold standard, two annotators worked independently to generate the datasets. If any discrepancies were found between the two annotators, they were reviewed among a third-party group to decide on the appropriate output, and the final gold standard datasets were produced.

### Evaluation of algorithms

The algorithms were evaluated using recall, precision, and F1-measure. Recall is the proportion of the gold standard entities that were correctly identified by the algorithm. Precision is the proportion of the extracted entities that were correctly found in the gold standard. The F1-measure is defined as 2*(precision*recall)/(precision+recall).

## RESULTS

### Performance

We evaluated each part of the algorithm separately. **Table 2a** presents the results for Part I on the test sets. The algorithm performed well for tacrolimus using both MedXN and medExtractR output; F1-measures were 1.00 for both NLP systems. For lamotrigine, the algorithm performed slightly better on the medExtractR output than the MedXN output. The medExtractR recall/precision/F1-measure was 1.00/1.00/1.00 while MedXN was 0.99/0.99/0.99.

**Table 2.** Recall, precision, and F1-measure performance on test sets.

| **a. Part I algorithm** |  |  |  |  |  |  |  |
| --- | --- | --- | --- | --- | --- | --- | --- |
|  |  | MedXN |  |  | medExtractR |  |  |
|  |  | Recall | Precision | F1 | Recall | Precision | F1 |
| Tacrolimus |  | 1.00 | 1.00 | 1.00 | 1.00 | 1.00 | 1.00 |
| Lamotrigine |  | 0.99 | 0.99 | 0.99 | 1.00 | 1.00 | 1.00 |
| **b. Part II algorithm (note/date)** |  |  |  |  |  |  |  |
|  |  | MedXN |  |  | MedExtractR |  |  |
|  |  | Recall | Precision | F1 | Recall | Precision | F1 |
| Dose Intake | Tacrolimus | 1.00/1.00 | 1.00/1.00 | 1.00/1.00 | 1.00/1.00 | 1.00/1.00 | 1.00/1.00 |
|  | Lamotrigine | 1.00/1.00 | 1.00/1.00 | 1.00/1.00 | 1.00/1.00 | 1.00/1.00 | 1.00/1.00 |
| Daily Dose | Tacrolimus | 1.00/0.99 | 1.00/1.00 | 1.00/0.99 | 1.00/0.99 | 1.00/1.00 | 1.00/0.99 |
|  | Lamotrigine | 0.98/0.97 | 0.98/0.97 | 0.98/0.97 | 0.99/0.99 | 0.99/0.98 | 0.99/0.98 |

The results for Part II can be found in **Table 2b**. Here we present the recall, precision, and F1-measure on the test sets for both medications at the note and date level, separated by the dose intake and daily dose. The note level collapsing performed very well with F1-measures of 1.00 for dose intake for both medications and both NLP systems, and F1-measures ranging from 0.98 to 1.00 for daily dose. Date level collapsing also performed well with F1-measures of 1.00 for dose intake and ranging from 0.97 to 0.99 for daily dose. The algorithm performed slightly better on dose intake than daily dose for both medications and both NLP systems.

### Error analyses and examples of challenges

Occasionally, the method for cost calculation we used in Part I did not result in the correct path being chosen, where a correct path is the path identified in the gold standard. **Table 3** shows an example of three competing paths and the cost calculations from each method using medExtractR output. In this example, all methods including our default method (minEntEnd) failed to choose the correct path, Path 1, as the entities were far apart from each other. The note read “100 mg tablet (Also Known As Lamictal);3.5 (three and one half) tablets by mouth twice a day Dispense: 210 tablets.” Our gold standard paired the dose amount “3.5” with the frequency “twice a day” while the algorithm with our default method paired the incorrectly extracted dose amount of “210” with the frequency of “twice a day”. In this case, the distance between “3.5” and “twice a day,” which is the correct pairing, is large enough that its cost is larger than that of the incorrect pairing of “210” with “twice a day,” despite a penalty the algorithm imposed when pairing a dose amount with a preceding frequency, which is unusual and often incorrect. An incorrect path could also be chosen when the greedy algorithm is used. For details about penalties and the greedy algorithm, see **Supplementary Material**.

**Table 3.** Example of incorrect path being chosen.

| **Standardized form:** |  |  |  |  |  |
| --- | --- | --- | --- | --- | --- |
| Dose Amount |  |  | Frequency |  |  |
| 3.5::184::187'210::248::251 |  |  | twice a day::226::237 |  |  |
|  | **Competing Paths** |  |  |  | **Path Chosen** |
|  | **Path 1 (Correct Path):** | **Path 2:** | **Path 3:** | **...** |  |
|  | 3.5 – twice a day<br>210 – NA | 3.5 – NA<br>NA – twice a day<br>210 – NA | 3.5 – NA<br>210 – twice a day |  |  |
| **Method** |  |  |  |  |  |
| byBeg | 37 | 32 | 35 |  | Path 2 |
| minEntBeg | 74 | 64 | 70 |  | Path 2 |
| sumBeg | 37 | 32 | 35 |  | Path 2 |
| byEnd | 35.5 | 32 | 29.5 |  | Path 3 |
| *minEntEnd\** | 71 | 64 | 59 |  | Path 3 |
| sumEnd | 35.5 | 32 | 29.5 |  | Path 3 |
| minMax | 60 | 69.7 | 46 |  | Path 3 |

\* The default method

The route entity occasionally caused discrepancies between the gold standard and the algorithm in Part I. The algorithm uses a standardized form of route, which sometimes leads to loss of information. For instance, a strength of 100 with routes of “po” and “oral” would only have a single row of output from the algorithm (since both expressions are standardized to “orally”) but two rows in the gold standard. Also, when making the gold standard, we did not allow strength to be borrowed across different standardized routes (e.g., orally and IV), but the algorithm did allow borrowing of strength in these cases.

In Part II, the intake time entity occasionally caused problems. When collapsing rows, intake time is only used to identify redundancies when part of a dose sequence. In cases where a lone am, noon, or pm row occurs, these rows would sometimes get removed as duplicates of other rows for the same note or date, despite one having an am/noon/pm and the other not having any intake time. Also, challenges occurred when the morning dose was in the row after the evening dose. The algorithm only treated dose sequences correctly when the morning dose came before the evening dose.

Another challenge occurred when the NLP extracted incorrect or incomplete information. An example of incorrect extraction can be seen in the example in **Table 3** where “210” was the number of tablets the pharmacy was supposed to dispense, but it was extracted as a dose amount. Incorrect extraction could also occur because of a misspelling, missing spaces, or an uncommon abbreviation of a drug name, which would cause the NLP to extract entities anchored to the wrong drug mention. This results in several extra strengths, dose amounts, or frequencies, making it difficult to get correct pairings. An example of an error due to incomplete extraction was observed for medExtractR (**Table 4**). Since we have a penalty in cases where a dose amount occurs after an intake time or frequency, the algorithm does not pair the “1” and “dinner” in this example. The gold standard, on the other hand, does pair these entities. Looking at the original note, we see the medExtractR output fails to capture some information (including “one half” and the second “dinner”), leading to difficulty in pairing the extracted entities. In general, when the NLP system extracts dosing information incorrectly, it is difficult for the algorithm to pair entities in a desirable manner (and usually does not produce the correct dose quantities based on the original clinical note).

**Table 4.**
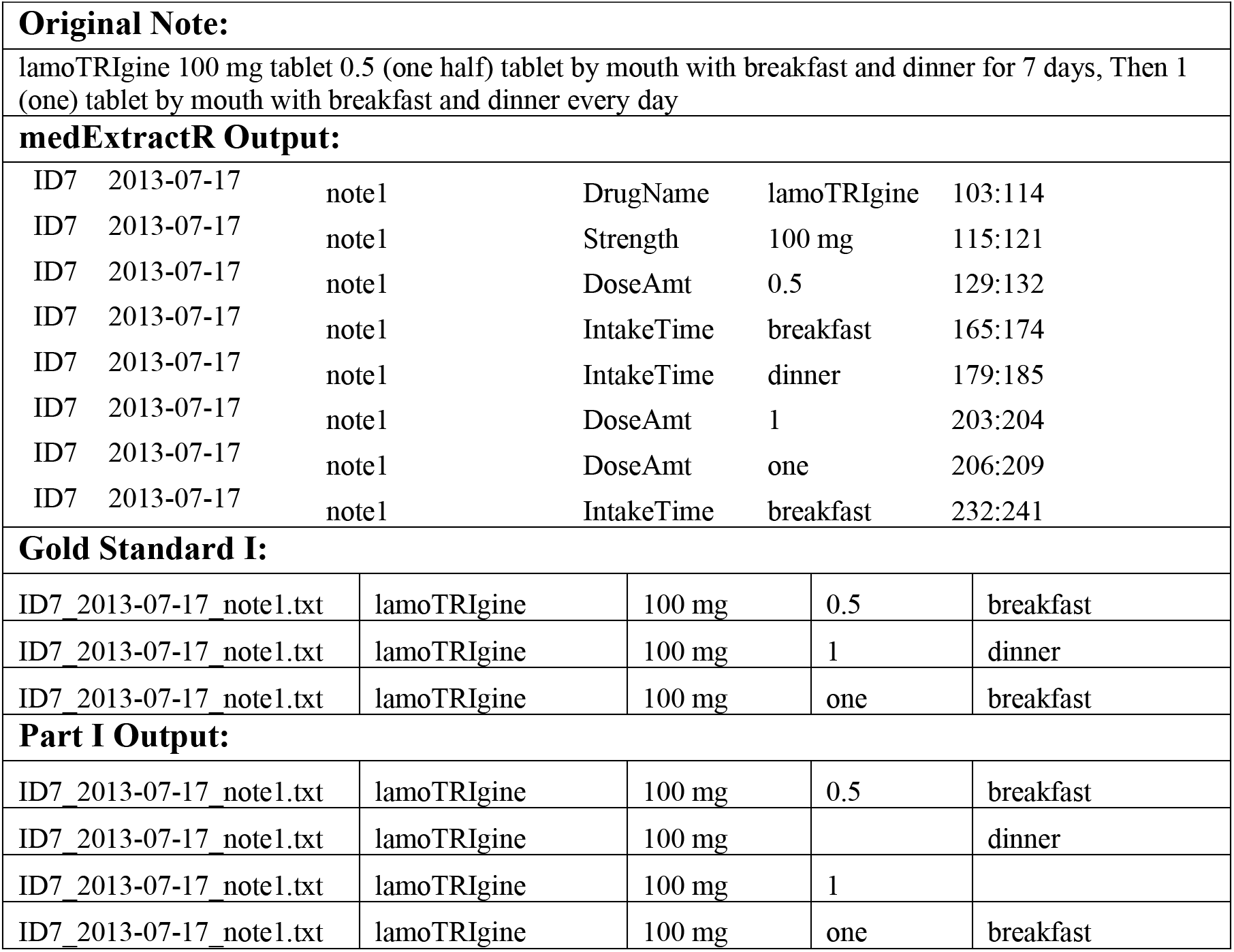
Example of medExtractR extracting incomplete information causing difficulty in pairing entities.

## DISCUSSION

Detailed medication dose information is often required to perform medication-based population studies. Medication dose data can be obtained from a structured data source or an unstructured data source such as clinical notes in EHRs. To extract medication dose information from unstructured text, a specialized algorithm such as an NLP system is commonly used. However, the output of NLP systems is often not in a form that is useful for analysis. In order to address this issue, we developed a post-processing algorithm. Our post-processing algorithm was developed using the output from two NLP systems, MedXN and medExtractR, which performed best at extracting medication information out of four NLP systems previously tested.[8] Our algorithm performed well to process and compute dose quantities from the output from both MedXN (F-measures Part I: ≥ 0.99; Part II: ≥ 0.97) and medExtractR (F-measures Part I: ≥ 1.00; Part II: ≥ 0.98).

Our algorithm has unique features. It was designed to have two distinct parts that were developed separately. Because the parts are separate, each can be easily modified if needed, while only minimally affecting the other part. This feature would also help the algorithm to be more generalizable to other NLP systems. Once the raw output from other NLP systems (e.g., MedEx or CLAMP) is converted to the standardized form (Step 1 of Part I), the algorithm is generally able to process it and generate the desired dose data. Some other NLP systems have a built-in step to pair dose entities, in which case only Part II of our algorithm would need to be applied. However, code for converting the raw output to the standardized form should be written by users who want to use different NLP systems. Though painstaking, as long as a gold standard can be generated, our algorithm can be tested on any NLP system.

We ran the algorithm on both the training and test sets using all of the cost calculation methods we considered and compared the precision and recall. We found that the minEntEnd and minEntBeg methods performed best and were very similar. Although we chose to make the minEntEnd method the default, we suggest that users run the algorithm with both methods. Identifying discrepancies between the methods can help users find potential issues with the raw NLP output.

There are some limitations in this study. First of all, though we tested the algorithm using two medications that have widely different prescribing patterns, it should be tested using other medications. Similarly, patterns which were common in Vanderbilt’s EHR system (e.g., “AKA” formats such as “tacrolimus 1 mg (also known as Prograf) 3 tablets bid”) may be represented differently at other institutions. Secondly, as shown in the error analyses, the algorithm cannot correctly process some cases. The default method we used for calculating cost in Part I did not always choose the correct path, especially when entities were far apart from each other. In Part II, rows were sometimes incorrectly removed as redundancies due to the way the algorithm treated dose sequences. Our error analyses suggest some improvement of the algorithm as future work.

The goal of the post-processing algorithm was to convert all the extracted information from an NLP system into a usable format for dose-related studies. However, this may result in conflicting doses on the same day, which will need to be resolved before the data can be used for medication-based studies such as pharmacokinetic or pharmacodynamic studies. Future work will include identifying the most likely correct dose when conflicting doses are present.

## FUNDING

This work was supported by the National Institute of General Medical Sciences under award R01-GM123109 (PI: Dr. Choi).

## AUTHOR CONTRIBUTIONS

As the principal investigator, LC conceived the research and managed the project. EM drafted the manuscript. CB implemented the post-processing algorithm using the R programming language. EM, HLW, MLW, and NTJ generated gold standard datasets and provided feedback to improve the post-processing algorithm throughout development. All authors reviewed and edited the final manuscript.

## CONFLICT OF INTEREST STATEMENT

None declared.

## Supplementary Material

### Supplementary Text

#### Cost calculation methods

For each of the possible paths in Step 4 of the Part I algorithm, we calculated the cost using the distance between entities. We considered seven methods for calculating cost. Each of the methods are described in **Table S1** and illustrated using an example in **Figure S1** of Supplementary Material. The distance between two entities is the number of characters between their start or end positions, depending on the cost method. In our examples, we consider the distances between dose amount, frequency, and duration, though the algorithm could apply to additional entities. When an entity is missing (i.e., NA), the distance is fixed at 32 as default, which can be changed by the user. We performed a sensitivity analysis to examine how different NA penalties would affect the results. We used penalties of 24, 26, 28, 30, 32, and 40. For penalties below 28, the recall and precision were worse for lamotrigine when using the medExtractR output. NA penalties of 28 and above did not change the performance for either drug or either NLP system. Based on the sensitivity analyses, we recommend using a penalty of 32 as the default.

When pairing dose amount and frequency, we have found that it is often undesirable for the dose amount to occur after the frequency. Hence, a penalty is added to the distance if the frequency occurs ahead of dose amount. This negative penalty should be supplied as a fraction (between 0 and 1) of the NA penalty. A penalty of 1 should ensure that dose amount is never paired with a preceding frequency, while the current default of 0.5 will allow it.

We tested all of the methods and found that the minimum distance between entity begin positions (minEntBeg) and the minimum distance between entity end position and next entity begin position (minEntEnd) methods performed best. We set the default method to be minEntEnd. **Figure S2** illustrates the cost calculation for each possible path using an example with the default cost method, minEntEnd. In this case, Path 1 has the minimum cost, so it would be selected as the proper pairing.

#### Greedy algorithm

When the cost calculation is performed for complicated cases that yield many combinations, the number of paths grows quickly. For these cases, finding the path with minimum cost can be computationally intensive. A greedy algorithm is applied for cases where the number of paths available for selection is deemed too large. This is based on an estimate of “n choose k”, where “n” is the number of pairs and “k” is the expected number of pairs to select. When this estimate exceeds a threshold, the greedy algorithm is used to select the approximate path rather than the optimal path. The default threshold is set at 100,000,000, which can be set by users.

---

**Supplementary Table S1.** Descriptions of distance method calculations. Define a segment to be the distance between entities in a pairing. Segments are formed between all entities in a pairing.

|  |  |
| --- | --- |
| **byBeg** | Calculate the length of each segment using the start positions of the entities. Find the sum of these lengths for each pairing in the path. The mean of these sums is the cost for the path. |
| **minEntBeg** | Calculate the length of each segment using the start positions of the entities. Find the minimum length from each possible pair of segments within each pairing in the path. Add these minimum lengths together for each pairing. The mean of these sums is the cost for the path. |
| **byEnd** | Calculate the length of each segment using the end position of the first entity and the start position of the second entity. Find the sum of these lengths for each pairing in the path. The mean of these sums is the cost for the path. |
| **minEntEnd** | Calculate the length of each segment using the end position of the first entity and the start position of the second entity. Find the minimum length from each possible pair of segments within each pairing in the path. Add these minimum lengths together for each pairing. The mean of these sums is the cost for the path. |
| **sumBeg** | Calculate the length of each segment using the start positions of the entities. Line up the entities in order by position for each pairing. Add together the length of the segments between the first and second entities and between the second and third entities in each pairing. The mean of these sums is the cost for the path. |
| **sumEnd** | Calculate the length of each segment using the end position of the first entity and the start position of the second entity. Line up the entities in order by position for each pairing. Add together the length of the segments between the first and second entities and between the second and third entities in each pairing. The mean of these sums is the cost for the path. |
| **minMax** | Line up the entities in order by position for each pairing. Find the distance between the start position of the first entity and the end position of the last entity for each pairing. If an entity is missing, add a penalty of 64 to the distance for that pairing. The mean of the distances is the cost for the path. |

**Supplementary Figure S1:**
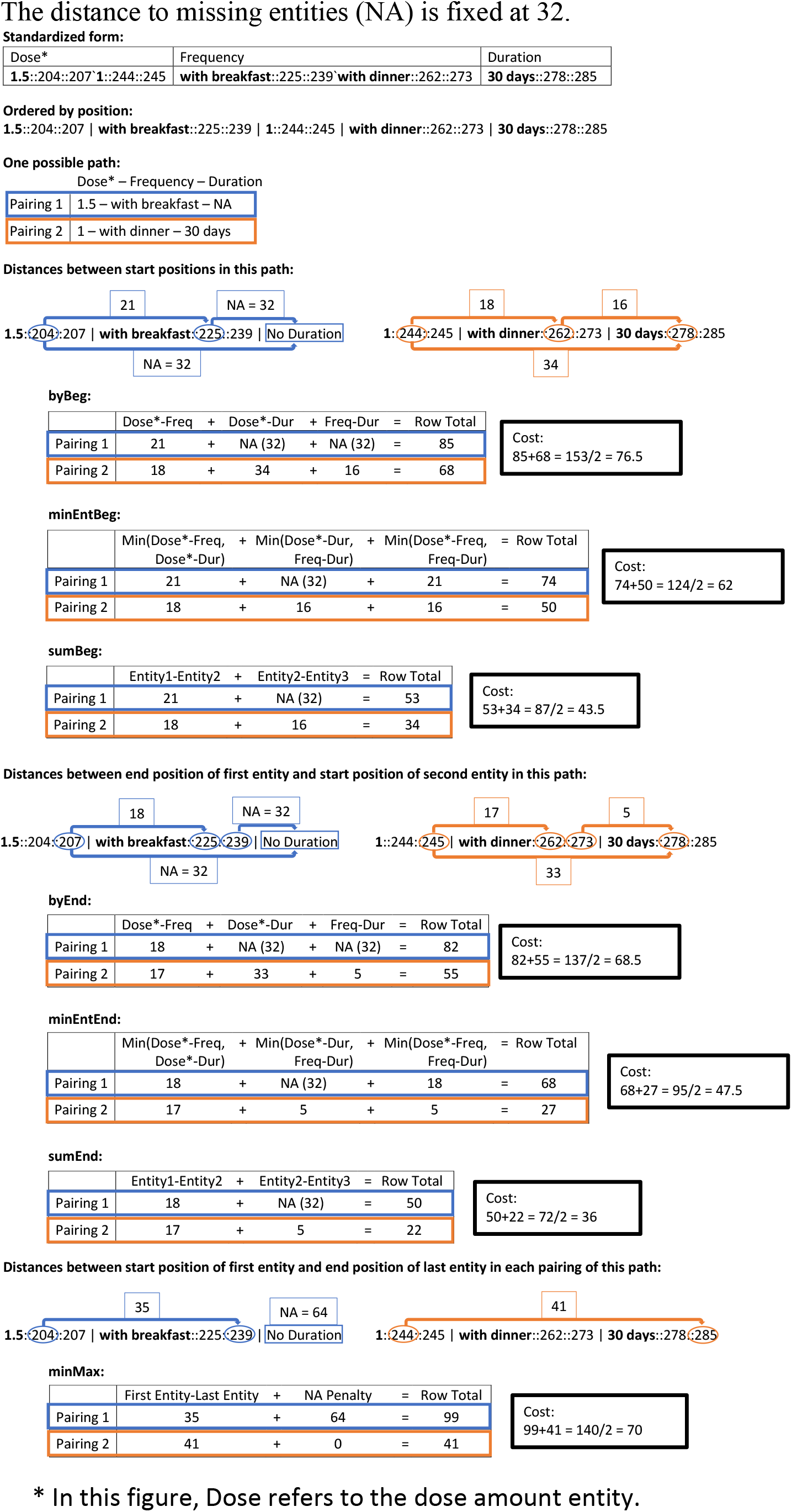
Examples of cost calculation methods. The distance to missing entities (NA) is fixed at 32.

**Supplementary Figure S2.**
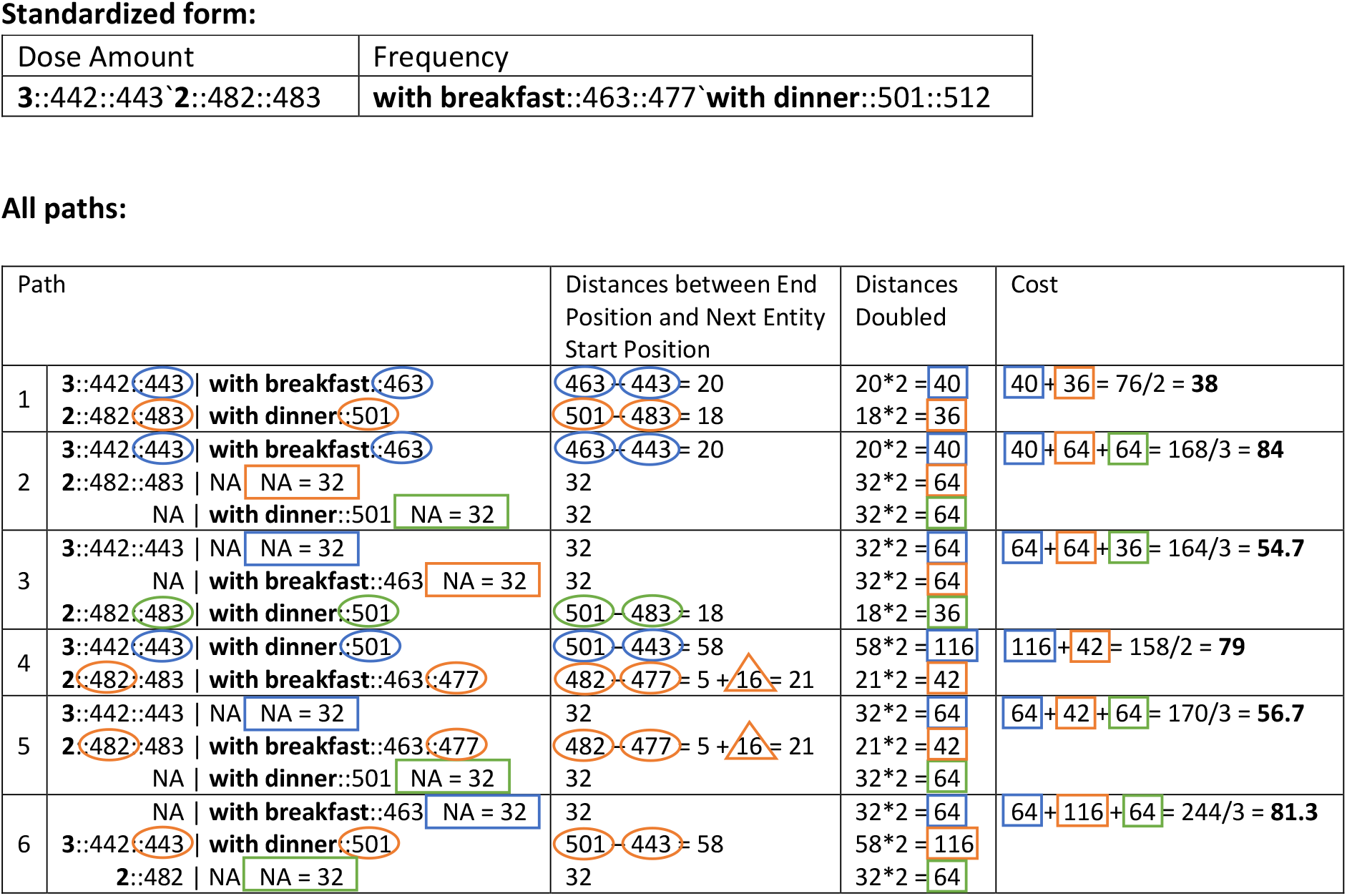
Example of cost calculation for all possible paths using the minEntEnd method. The distance to missing entities (NA) is fixed at 32. Path 1 has the minimum cost of 38, so it would be selected. Circles, rectangles, and triangles represent end/start positions, the fixed distance of 32, and the penalty for frequency occurring before dose amount, respectively. Each color represents a different pairing. For the minEntEnd method, distances are doubled when there are only two entities.

